# QTL spanning the TGF-β2 locus is associated with muscle fiber hypertrophy in rainbow trout

**DOI:** 10.64898/2026.05.24.727516

**Authors:** Akash Raghu, Guglielmo Raymo, Ridwan Ahmed, Ali Ali, Tim Leeds, Mohamed Salem

## Abstract

**Background:** Skeletal muscle growth is a key determinant of body size and market value in salmonid aquaculture, yet the mechanisms linking genomic variation to muscle fiber hypertrophy remain poorly resolved. Myofiber cross-sectional area (CSA) provides a quantitative cellular proxy for fiber size and a direct link to macroscopic growth traits.

**Methods:** We performed histological phenotyping of white skeletal muscle from rainbow trout (*Oncorhynchus mykiss)* representing divergent fillet-yield selection lines (ARS-FY-H and ARS-FY-L), quantifying mean myofiber CSA and fiber number using high-throughput image analysis. Genome-wide association analysis (GWAS) was conducted using low-pass whole-genome sequencing (∼1×) with genotype imputation and functional variant annotation. RNA sequencing was performed on fish representing high and low CSA extremes to identify differentially expressed genes and enriched biological pathways.

**Results:** Mean myofiber CSA was significantly associated with body weight, muscle weight, visceral weight, and body length (p < 0.05), while fiber count showed no significant association with most growth traits, implicating hypertrophy as the primary driver of muscle mass variation. GWAS identified a significant QTL spanning ∼4.76 Mb on chromosome 2 (117 significant SNPs; Bonferroni-adjusted P ≤ 0.05; λ = 1.02). Associated variants were predominantly noncoding, enriched in intronic, intergenic, and enhancer-annotated regions. A high density of SNPs colocalized with the TGF-β2 locus, overlapping strong and genic enhancer elements in white muscle. Transcriptomic comparisons revealed that high-CSA muscle showed elevated expression of genes related to contractile function, cytoskeletal organization, and translation, while low-CSA muscle exhibited upregulation of extracellular matrix and immune-related genes consistent with a tissue remodeling state.

**Conclusions:** Noncoding regulatory variation within a significant QTL spanning the TGF-β2 locus is associated with distinct transcriptional programs linked to muscle fiber hypertrophy in rainbow trout. By integrating genetic variation, chromatin-state annotation, and transcriptomic profiling, this study identifies candidate regulatory loci associated with variation in muscle cellularity and growth-related phenotypes in rainbow trout.

## BACKGROUND

Skeletal muscle growth is a primary determinant of vertebrate body size and a key complex trait shaped by interactions between genetic variation and cellular processes. In salmonids, enhanced muscle growth improves feed conversion efficiency and increases market value per fish, and has been a major target of selective breeding programs. To this end, the USDA’s National Center for Cool and Cold Water Aquaculture (NCCCWA) has implemented long-term selective breeding programs to increase muscle yield and body weight in rainbow trout (Oncorhynchus mykiss) populations [1].

Post-embryonic muscle growth in teleost fish is a plastic process driven by hypertrophy (increase in fiber size) and hyperplasia (recruitment of new fibers) [2–4]. While hyperplasia contributes substantially to early growth, hypertrophy becomes an increasingly dominant driver of muscle mass as fish increase in body length [5–7]. Mean myofiber cross-sectional area (CSA), defined as the transverse area of individual muscle fibers, provides a quantitative proxy for fiber size and a cellular link to organismal growth traits. This metric provides a robust cellular correlate of muscle growth, linking microscopic tissue architecture to macroscopic phenotypes including body weight, length, and muscle mass. Histological quantification of muscle fiber CSA is widely used to assess muscle cellularity and growth dynamics in teleost fish [6, 8].

Despite the commercial importance of muscle mass in salmonids, the mechanisms linking genomic variation to muscle cellular phenotypes remain incompletely understood. Current research suggests that high-yield phenotypes are driven by a combination of protein accretion and reduced protein degradation processes [1, 9, 10]. However, the genetic architecture underlying variation in muscle fiber size, particularly the contribution of regulatory variation, remains unclear.

Genome-wide association studies have identified loci associated with growth traits in salmonids; however, these signals often map to broad genomic regions and are enriched for noncoding variants, limiting mechanistic interpretation. In particular, the extent to which regulatory variation contributes to differences in muscle fiber hypertrophy remains poorly resolved.

In vertebrates, muscle growth is regulated by conserved pathways such as IGF-1/mTOR signaling, myostatin inhibition, and satellite cell proliferation, which coordinate protein synthesis, fiber hypertrophy, and regeneration [8]. While these pathways provide a functional framework, their genomic regulation and relative contributions to muscle fiber hypertrophy in salmonids remain unclear.

The objective of this study was to characterize the genetic and transcriptomic basis of variation in muscle fiber size in selectively bred rainbow trout. Here, we combined histological phenotyping with high-throughput image analysis to quantify muscle fiber size and density. Transcriptome profiling was performed to identify gene expression differences associated with muscle cellular phenotypes. We further utilized a Genome-Wide Association Study (GWAS) to map these traits to the rainbow trout genome, identifying a significant quantitative trait locus (QTL) on chromosome 2. This QTL spans 117 significant SNPs, including a high concentration within the *TGF-β2* (Transforming Growth Factor Beta-2 Proprotein) gene, a known regulator of cellular proliferation and differentiation. By integrating histological, genomic, and transcriptomic data, this study provides a framework linking genetic variation to muscle cellularity and gene expression, and highlights the role of regulatory variation in shaping muscle fiber hypertrophy.

## MATERIALS AND METHODS

### Rainbow trout population, experimental design, and sampling

This study utilized rainbow (*Oncorhynchus mykiss)* from the muscle-yield selection lines developed at the USDA National Center for Cool and Cold Water Aquaculture (NCCCWA). The base population was established as a growth-selected line in 2002 and underwent five generations of selection for improved growth performance [11]. Subsequent generations were selected for fillet yield following the protocols described by [1]. Fish from the 2020 year class (YC) were used in the present study, representing third-generation families from lines divergently selected for high (ARS-FY-H) or low (ARS-FY-L) fillet yield, as previously described [12].

### Family structure, rearing conditions, harvest, and tissue collection

Immature, all-female fish originated from 40 full-sib families (20 families per line; ARS-FY-H and ARS-FY-L) and were transferred from the NCCCWA to the University of Maryland, College Park (Crane Aquaculture Facility) at 322 days post-hatch. Fish were reared in a recirculating aquaculture system (RAS) with routine monitoring of water quality parameters. For approximately six months prior to harvest, fish were fed an astaxanthin-supplemented commercial diet (BioTrout pellets, 4.0 mm and 6.0 mm; ∼40 ppm astaxanthin; Bio-Oregon). At 442–485 days post-hatch, a total of 442 fish were harvested (mean body weight = 694.36 g; SD = 173.76 g).

Fish were withheld from feed for 24 h prior to harvest and were humanely euthanized. Phenotypic data, including total body weight (g), fork length (mm), and visceral weight (g), were recorded for each individual. Muscle weight (g) was determined by weighing the trimmed, skinless fillets. For histological and molecular analysis, white muscle tissue samples were collected from the epaxial region directly ventral to the dorsal fin. Muscle yield was expressed as a percentage of total body weight, calculated by the ratio of muscle weight to body mass as previously described [12, 13].

### Histological Phenotyping

Histological phenotypes were generated from a subset of 85 fish originating from 19 distinct genetic families (genetic lines and phenotypic traits of individual fish are provided in Supplementary File 1). Fish were selected to maximize representation across families and variation in growth-related phenotypes while maintaining balanced representation of the FY-H and FY-L selection lines. Of the selected fish, 42 belonged to the high fillet-yield genetic line (ARS-FY-H) and 43 belonged to the low fillet-yield genetic line (ARS-FY-L). Specifically, white muscle samples collected under the dorsal fin were fixed in Bouin’s solution for 24 h and then transferred to 70% ethanol for long-term storage. Tissues were trimmed, dehydrated, and embedded in paraffin wax. Sectioning was performed using Superfrost Plus slides (Thermo Fisher Scientific, Waltham, MA, USA). Serial sectioning was conducted, producing 4 μm thick slices with a 200 μm gap between sections to ensure representative sampling.

### Image Acquisition and Statistical Analysis

Histological sections were examined at 20× magnification using a Zeiss Axioplan EL-EINSATZ (451888) Phase Contrast Microscope and Zeiss AxioCam MRc 1.4MP Microscope Camera. For each fish, three non-overlapping fields were imaged per slide from two slides (six images total). Images were analyzed in MIPAR Image Analysis Software (v4.3.0) using a custom workflow for myofiber segmentation and background removal to quantify myofiber number and CSA (Figure 1).

**Figure 1.**
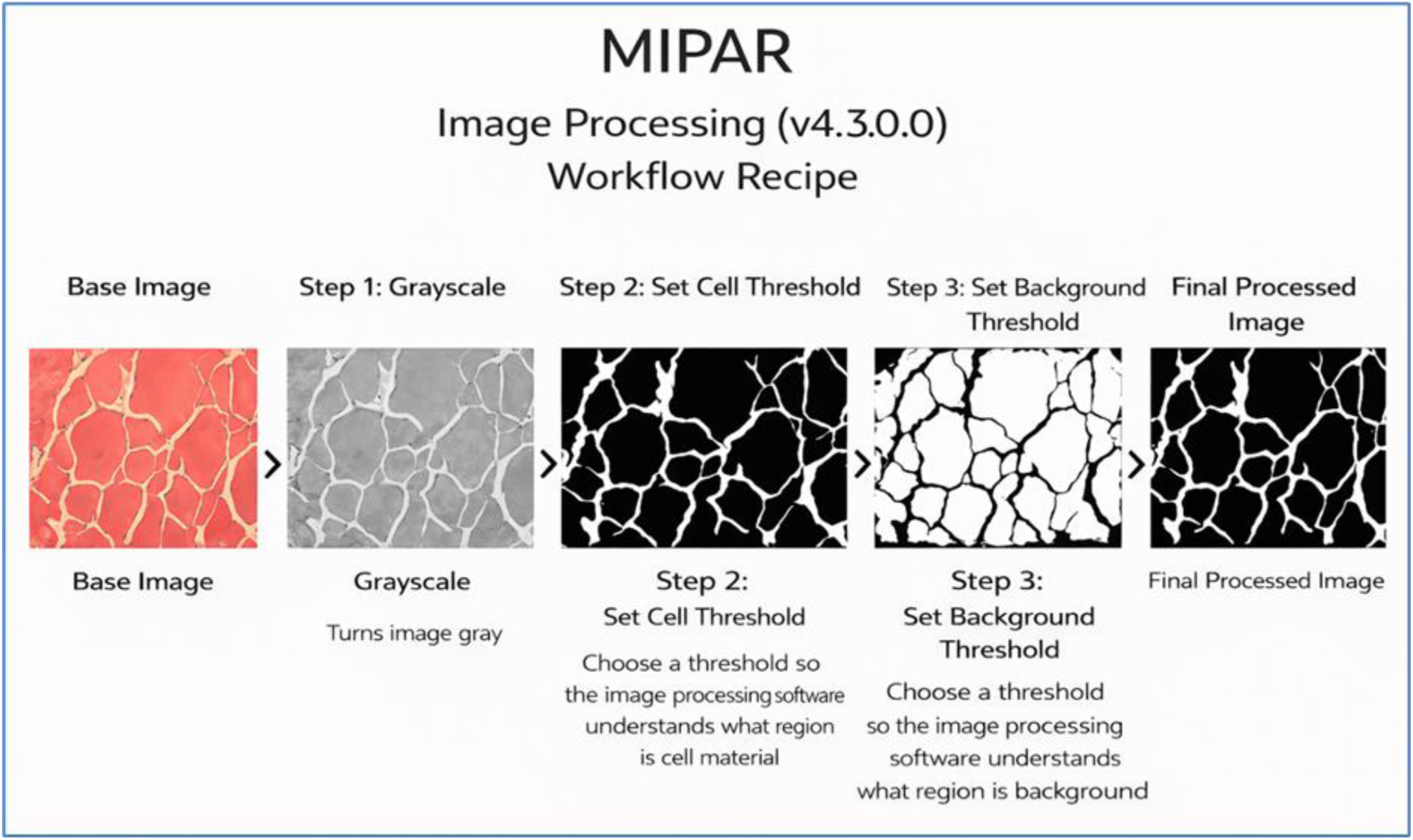
MIPAR image processing workflow (v4.3.0) used to derive myofiber number and CSA from muscle histology images. The pipeline shows the base image, grayscale conversion, cell thresholding, background thresholding, and background subtraction to generate the final processed image used for quantification.

Myofiber size was quantified as mean CSA (µm²) measured on transverse sections. Fibers smaller than 176.8 µm² (converted from the original image-analysis threshold of 1,700 px²; 1 px² = 0.104 µm²) were excluded to reduce background noise. This cutoff was determined empirically by testing multiple size thresholds; inclusion of objects below 176.8 µm² consistently introduced small artifacts and segmentation noise, whereas thresholds at or above 176.8 µm² produced the cleanest images with well-defined fibers. Results were qualitatively stable across nearby thresholds, indicating that downstream conclusions were not sensitive to minor adjustments of this cutoff. The workflow was run in batch using the MIPAR Batch Processor (v4.3.0).

Fields of view were selected from regions containing clearly delineated, intact fibers and free of obvious artifacts (e.g., tissue folds, debris, or poorly segmented areas). Within these quality-controlled regions, image capture was randomized across each sample to avoid systematic regional bias. For each fish, CSA and fiber count were averaged across the six images to generate one value per sample (n = 85). Fiber count did not differ between groups and was not included in downstream phenotypic correlations. Since total image area also includes variable amounts of interstitial and extracellular space surrounding fibers, this can confound the purported near-perfect negative relationship between filled area and fiber count.

Associations between myofiber CSA and growth traits were assessed using simple linear regression; model fit is reported as R² and significance was evaluated using two-sided tests (α = 0.05). Fish with higher myofiber CSA showed greater body length, body weight, visceral weight, and muscle weight. Based on these relationships, fish representing the upper and lower extremes of myofiber CSA were selected for RNA sequencing.

### DNA Extraction, Whole-Genome Sequencing, and Genotype Imputation

Genomic DNA was isolated from livers originating from individual fish (n = 85) using a modified salting-out protocol [14], with quality and quantity verified by gel electrophoresis and Qubit fluorometry. Paired-end libraries (insert size ∼350 bp) were prepared with the Illumina Nextera DNA Flex kit and sequenced on an Illumina NovaSeq 6000 platform (2 × 150 bp), generating low-pass whole-genome coverage of approximately 1× per sample. Genotypes were inferred using Gencove’s imputation workflow (Gencove, Inc.), which employs a modified implementation of GLIMPSE2 [15] and a haplotype reference panel built from 10×–30× whole-genome sequences of 160 diverse wild and domesticated rainbow trout (BioProjects PRJNA386519 and PRJNA402066) aligned to the USDA_OmykA_1.1 reference genome. Leave-one-out cross-validation yielded a mean genotype concordance of 96.5% for high-confidence sites (posterior probability > 0.90).

### Genome-Wide Association Study (GWAS) and SNP Annotation

Following imputation, variants with genotype probabilities below 0.90 were removed. Additional quality control was performed using PLINK v1.90p [16]. Variants were filtered based on a minor allele frequency (MAF) > 0.10, genotype missingness ≤ 0.0005, and Hardy–Weinberg equilibrium (HWE) exact test P > 0.10. After quality control, a total of 87,897 SNPs across 85 samples were retained for downstream quantitative trait analyses. Genome-wide association analysis was performed using a linear model in PLINK v2.0, with mean myofiber CSA as the response variable and SNP genotype as the predictor, including the first 20 principal components (PCs) and family as covariates to account for population structure and relatedness. Principal components derived from principal component analysis (PCA) were included to reduce false-positive associations arising from population stratification and ancestry-related genetic structure. Family was included as an additional covariate to further account for shared genetic background among closely related individuals within the experimental population. Model performance was evaluated using genomic inflation analysis, which showed minimal inflation (λGC = 1.02; Supplementary Figure 1), indicating that population structure was adequately controlled and that the model was not substantially over-parameterized. Although myofiber CSA was significantly associated with body weight, the relationship was modest (R² = 0.087), indicating that variation in CSA was not explained solely by overall body size. To further evaluate whether the association signal reflected CSA-specific effects rather than general body size, GWAS analyses were repeated including body weight as an additional covariate. The significant association signal remained localized to chromosome 2, indicating that the identified QTL was robust to inclusion of body weight in the model.

Significant SNPs were functionally annotated with SnpEff v5.2 using a custom database generated with the USDA_OmykA_1.1 rainbow trout reference genome (GenBank: GCA_013265735.3), including protein-coding and CDS tracks [17]. Associated variants were then visualized in the IGV genome browser (v2.19), overlaid with gene annotations and chromatin state tracks from multiple rainbow trout tissues [18] to evaluate their potential regulatory and coding impacts.

### RNA Extraction

At harvest, muscle tissues were flash-frozen in liquid nitrogen and stored at −80°C until processing. Total RNA was extracted using RNAzol (Molecular Research Center Inc., USA) following the manufacturer’s protocol. RNA concentration was measured using a NanoDrop spectrophotometer (Gen5 v2.09.2; BioTek Instruments, Inc., USA), and purity was assessed using the A260:A280 ratio; samples with values of 1.8–2.1 were retained. RNA integrity was evaluated by gel electrophoresis and further assessed using the ScreenTape® system (Agilent, Santa Clara, CA). Only samples with RIN ≥ 7 were used for library preparation.

### RNA-Seq Library Preparation and Sequencing

Transcriptome profiling was performed on skeletal muscle from 11 fish selected from the upper (n = 5) and lower (n = 6) extremes of the mean myofiber CSA distribution. Samples were drawn from four families to reduce potential confounding from shared genetic background. To minimize genetic background effects, selections were restricted to two families per line.

Library preparation was conducted with the NEB polyA selection module (Ipswich, MA, USA) in the Integrated DNA Technologies (Coralville, IA, USA) xGen RNA library preparation kit. The preparation was performed according to the manufacturer guidelines. First RNA undergoes fragmentation, followed by random priming and reverse transcription to generate first-strand cDNA. Next tailing and adapter ligation to the 3’-end of the cDNA molecule is conducted and a final PCR is conducted to increase library yield. Sequencing was performed at the Oklahoma Medical Research Foundation NGS Core, USA using an Illumina NovaSeq -S4 Instrument and paired end 150 cycle sequencing.

### Differential Gene Expression Analyses

Initial quality control, including low-quality read removal, adapter trimming, and read mapping, were all performed using the CLC genomics workbench (version 22.0, Qiagen, Hillden, Germany). Processed reads were mapped to the rainbow trout reference genome (NCBI Accession GCF_013265735.2). Reads were mapped using a mismatch cost of 2. Raw counts were used to identify differentially expressed genes (DEGs) between high and low CSA individuals using DESeq2 (v1.46) with batch effect correction enabled [13]. The differential gene cutoff was determined by P-adjusted-value < 0.05.

Functional enrichment analysis was performed to identify KEGG (Kyoto Encyclopedia of Genes and Genomes) pathways [19] and GO (Gene Ontology) terms based on the DEGs identified against the background (expressed) genes in muscle. The analysis was performed using ShinyGO 0.741[20] with default setting. A cut off of FDR-adjusted P-values of less than 0.05 was used for significant GO terms and KEGG pathways.

## RESULTS AND DISCUSSION

We evaluated the relationship between muscle cellularity and growth traits using histological phenotyping of myofiber cross-sectional area (CSA) and fiber number in selectively bred rainbow trout. Using USDA-provided families representing divergent fillet-yield selection lines, we quantified mean myofiber CSA, a proxy for fiber size, and tested its association with body weight, muscle weight, visceral weight, muscle yield, and body length. Overall, our data show a significant, albeit modest, association between muscle cellularity, particularly fiber size, and growth-related phenotypes in rainbow trout. The ARS-FY-H and ARS-FY-L lines were included because they represent populations divergently selected for fillet yield across multiple generations, providing a biologically relevant framework to evaluate whether long-term selection for harvest-related traits is accompanied by differences in muscle cellularity and myofiber hypertrophy. In the 2020 year class used here, third-generation FY-H families exhibited greater mean body weight and fillet yield than FY-L families (1249 g vs. 1137 g body weight; 57.1% vs. 54.2% fillet yield), consistent with the expected phenotypic divergence between lines. Although the FY-H and FY-L lines differ in growth and fillet-yield traits at the population level, the present analyses focused primarily on variation in myofiber CSA across individuals rather than formal contrasts between selection lines. Furthermore, the high-, intermediate-, and low-CSA groups consisted of heterogeneous mixtures of families and genetic backgrounds, indicating that the observed associations are unlikely to reflect line effects alone.

Mean myofiber CSA was positively associated with multiple growth traits. Regression analyses showed significant, albeit modest, associations between CSA and body weight, muscle weight, visceral weight, and body length (p < 0.05), indicating that increased fiber size is associated with greater somatic growth in this population (Table 1). In contrast, fiber count was not significantly associated with most growth traits, suggesting that hypertrophy, rather than hyperplasia, may contribute more substantially to variation in muscle mass at this developmental stage. These findings are consistent with previous observations across vertebrates, where muscle fiber size is positively associated with growth-related traits, while fiber density often shows weaker or inverse relationships [21]. In teleosts, hypertrophy has been reported to become the dominant contributor to muscle mass accretion at later growth stages, supporting the interpretation that variation in myofiber CSA reflects differences in growth trajectory rather than muscle proportion alone [3, 8].

**Table 1.** Association between myofiber CSA and key performance traits. Mean myofiber CSA was significantly associated (P < 0.05) with higher body weight, muscle weight, visceral weight, and total body length.

| Fish Characteristic | Myofiber CSA R<sup>2</sup> | Myofiber CSA P-Value | Cell Count R<sup>2</sup> | Cell Count P-Value |
| --- | --- | --- | --- | --- |
| Body Weight | 0.087 | **0.007\*** | 0.016 | 0.252 |
| Muscle Weight | 0.079 | **0.010\*** | 0.008 | 0.436 |
| Visceral Weight | 0.071 | **0.017\*** | 0.05 | **0.044\*** |
| Muscle Yield | 0.005 | 0.421 | 0.02 | 0.216 |
| Body Length | 0.078 | **0.012\*** | 0.003 | 0.668 |

Notable size-dependent patterns of white muscle hypertrophy have been reported in other teleost species. In Nile tilapia, morphometric analyses across length classes indicate that hypertrophy becomes a primary contributor to increasing muscle mass as fish grow [22]. In rainbow trout, moderate sustained exercise can also shift white muscle fiber traits, illustrating that cellularity metrics are responsive to production-relevant physiological contexts [23]. In contrast, fiber count was not significantly associated with most growth traits, including muscle weight (p = 0.436). The only significant relationship for fiber count was with visceral weight (p = 0.044), suggesting that variation in fiber number may extrapolate to broader differences in body composition rather than strict muscle accretion.

The limited association between fiber count and most growth traits is consistent with salmonid work showing that fiber recruitment and fiber density can vary substantially among strains, year classes, and rearing environments, which can reduce simple linear relationships at a single sampling time point [2, 24]. Notably, in our data, muscle yield (%) was not significantly correlated with either myofiber size or fiber count, implying that hypertrophy increases absolute size but does not necessarily shift the relative proportion of muscle to total body mass.

Similarly, uncoupling between muscle cellularity and harvest-relevant outcomes has been reported in other cultured fish. In sea bass, muscle cellularity measures were evaluated alongside flesh quality traits at commercial size, indicating that microstructure contributes to product characteristics but is not a single determinant [25]. In Atlantic salmon, comparisons of wild and farmed fish likewise show that differences in texture and related quality traits can reflect multiple contributors beyond fiber cellularity, including connective tissue and post-rigor properties [26].

Figure 2 shows evident clustering of individuals into Bottom, Intermediate, and Top groups along the myofiber CSA axis, consistent with group assignment based on myofiber CSA ranking. Across all panels, myofiber CSA exhibited a consistent positive trend with body length, body weight, visceral weight, and muscle weight (see Table 1 for regression statistics). This relationship aligns with established teleost growth patterns in which increases in muscle fiber size scale with somatic growth; for example, muscle fiber diameter increases with body length during early development in European sea bass [27], and hypertrophy is widely recognized as a major contributor to muscle mass accretion as fish approach later growth stages [3]. Although the ranking groups cluster strongly along the x-axis by design, separation along the growth traits is incomplete, and substantial overlap among groups is observed for each phenotype. This overlap is expected because body size and tissue weights reflect multiple contributing processes, including developmental timing, tissue allocation, and body composition, rather than local muscle cellular area alone [3]. The positive trend with visceral weight further suggests that myofiber CSA may track broader growth and nutrient partitioning patterns rather than muscle accretion in isolation.

**Figure 2.**
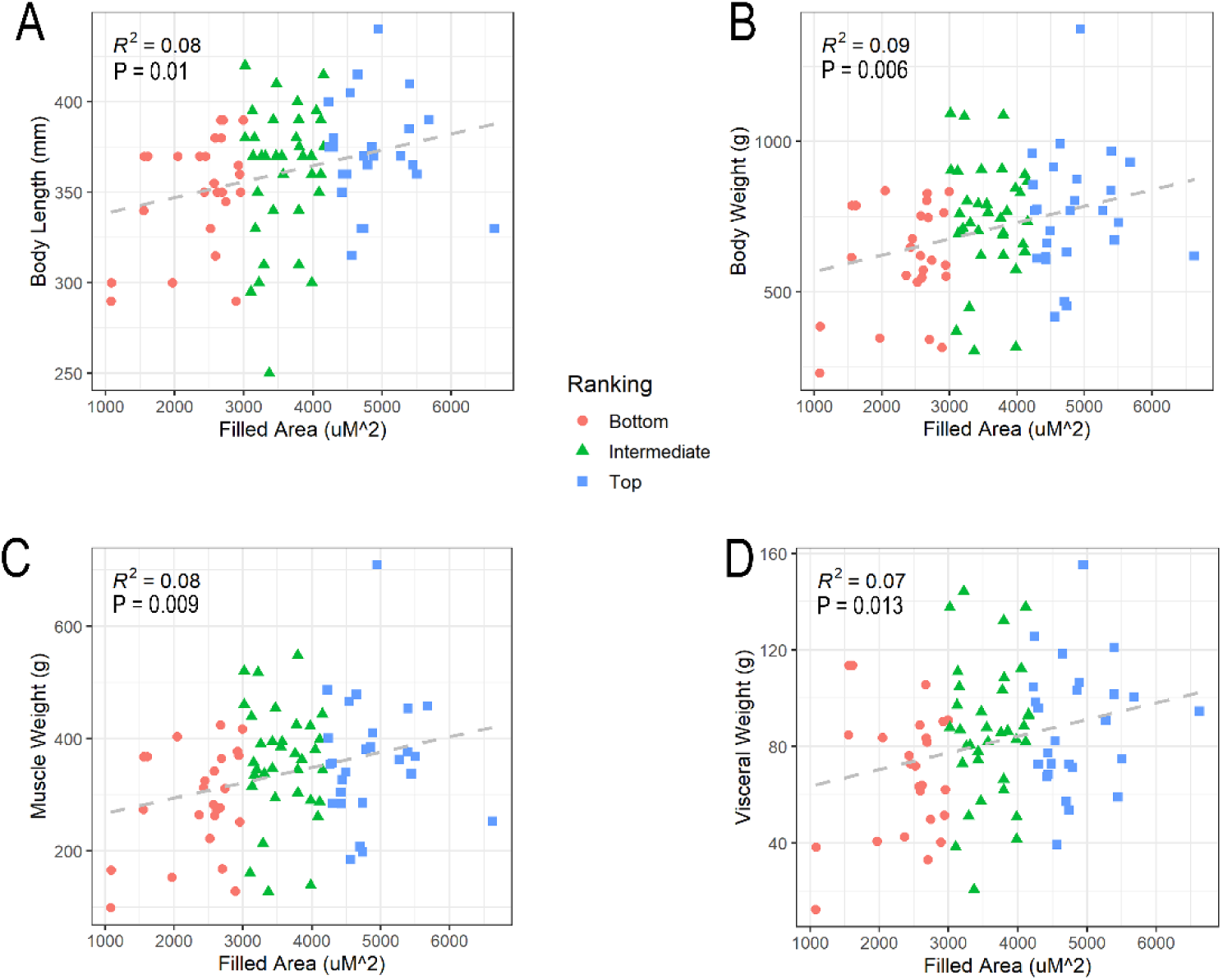
Distribution of myofiber CSA groups across phenotypic characteristics. The y-axis shows (A) body length (mm), (B) body weight (g), (C) visceral weight (g), and (D) muscle weight (g). Individuals in the top and bottom 25% of myofiber CSA are highlighted, illustrating variability in muscle fiber size across phenotypes. The x-axis denotes mean myofiber CSA for each sample reported as cross-sectional area (µm²) converted from MIPAR output values (1 px² = 0.104 µm²). Linear models accounting for genetic heterogeneity were fit for each trait, and corresponding adjusted R² and slope P-values for myofiber CSA are shown. Predicted adjusted regressions are denoted by dashed lines, with prediction confidence intervals shown in grey.

Building on the histological association between mean myofiber CSA and growth, we used GWAS to identify associated genetic variants and then applied RNA-seq to define the transcriptional programs distinguishing high- and low-myofiber CSA.

### Genome-Wide Association Study Reveals Notable QTL on Chromosome 2 Associated With Increased Myofiber CSA

Genome-wide association analysis identified 117 SNPs exceeding the genome-wide threshold applied in this study (Figure 3). Genomic inflation (λGC) was calculated to be 1.02, suggesting there is no bias or inflation in our analysis. All signals were localized to a single region on chromosome 2 spanning an ∼4.76 Mb interval (Figure 3a), consistent with a significant QTL associated with variation in myofiber CSA. Reanalysis of the GWAS including body weight as an additional covariate produced a similar association profile (data not shown), with significant SNPs remaining localized to chromosome 2. Importantly, the chromosome 2 association signal remained robust after inclusion of body weight as a covariate, indicating that the detected QTL was not attributable to overall body size and further supporting an association between the chromosome 2 locus and variation in myofiber CSA. Within this region, many SNPs exhibited near-identical P-values, consistent with strong linkage disequilibrium across the interval, such that multiple variants capture the same underlying association signal (Figure 3b-d). The predominance of noncoding variants within this region suggests that the association with myofiber CSA is primarily mediated through regulatory mechanisms rather than protein-coding changes.

**Figure 3.**
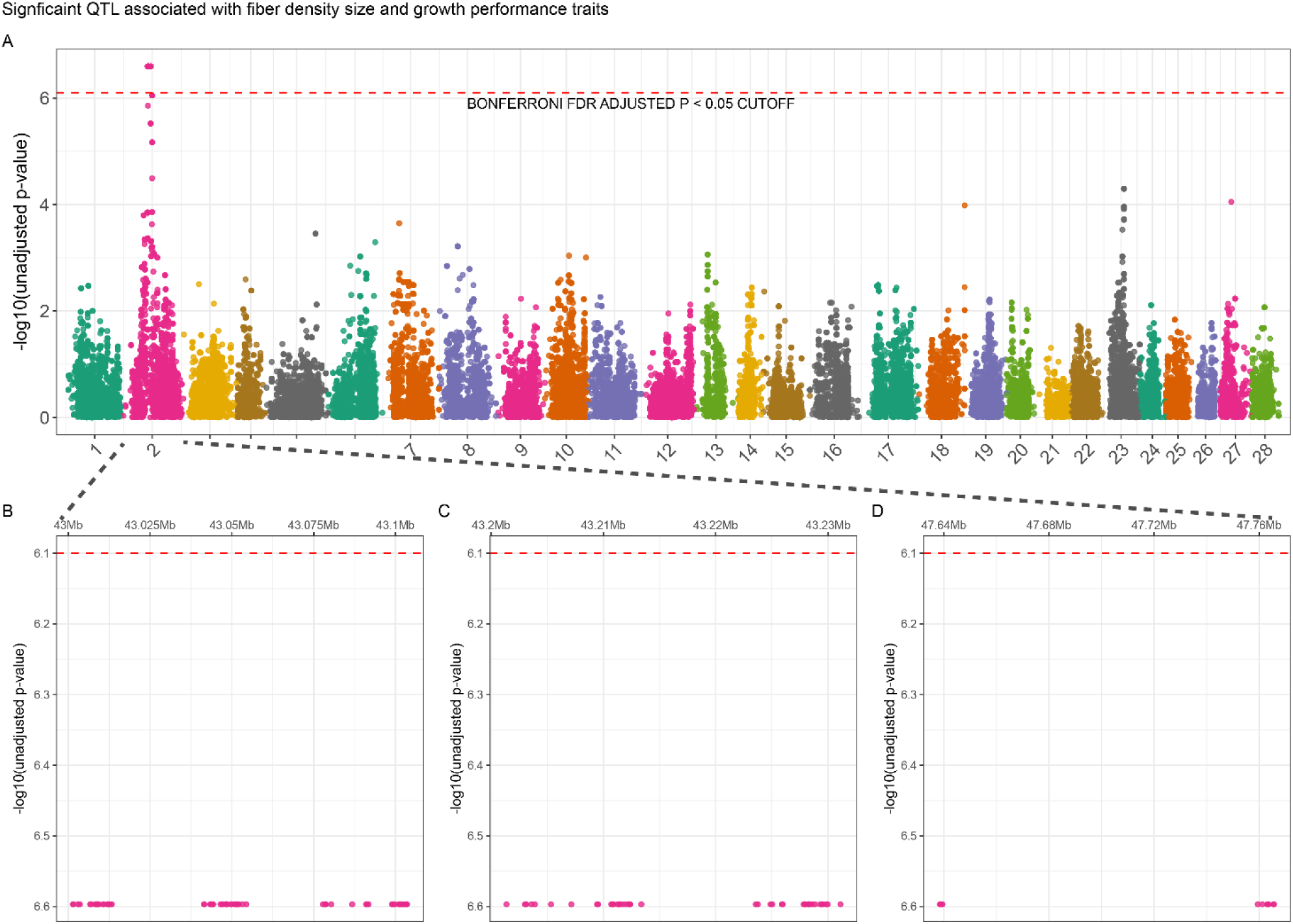
(a) Manhattan plot depicting SNPs distributed across autosomes. Horizontal dashed line denotes statistical significance (Bonferroni-adjusted P ≤ 0.05) when assessing myofiber CSA as a linear variable. (b-d) Genomic windows on chromosome 2 harboring SNPs exceeding the statistical significance threshold depicted by horizontal dashed line.

The breadth of the chromosome 2 QTL interval likely reflects extended linkage disequilibrium within this structured pedigree rather than a single causal locus, a recognized limitation in closed breeding populations with small effective population sizes. Fine-mapping through haplotype-based tests, conditional analyses, and replication across independent year-classes will be needed to narrow the signal and prioritize causal variants prior to application in selection. The candidate genes identified here, particularly *TGF-β2*, should therefore be considered priorities for functional validation pending such refinement.

Several genes within the chromosome 2 QTL provide biological context for muscle phenotype variation. A high density of associated SNPs mapped to *TGF-β2,* including intronic and putative regulatory annotations (Table 3). Notably, *TGF-β* signaling can inhibit myogenic differentiation and regeneration and modulate remodeling processes in skeletal muscle, including effects on myoblast differentiation/fusion and tissue repair dynamics [28–30]. In rainbow trout specifically, TGF-β2 shows dynamic expression during muscle growth resumption and satellite cell differentiation, supporting its relevance to muscle growth regulation in this species [31]. TGF-β2 and myostatin both belong to the broader TGF-β superfamily; however, they generally signal through distinct receptor pathways. In trout, strong muscle-mass effects have been demonstrated for the activin type IIB receptor pathway, which mediates signaling by myostatin and other activin-family ligands, highlighting that multiple branches of the TGF-β superfamily can influence muscle growth [32].

The chromosome 2 QTL also overlapped with candidate genes with noted translational capacity and cellular organization potential. Namely, *RRP15* functions in pre-rRNA processing and ribosome biogenesis, two processes directly linked to protein synthesis capacity [33]. *CNIH4* has been described as a cargo-sorting receptor that recruits specific client proteins into COPII vesicles, supporting ER export and ER-to-Golgi trafficking [34]. *FEZ2* belongs to the *FEZ/UNC-76* family implicated in intracellular transport and cytoskeletal-associated trafficking in other vertebrate systems, providing additional supporting context for this QTL and its potential influence on muscle cellular architecture and proliferation [35].

Collectively, the chromosome 2 QTL highlights candidate mechanisms spanning growth-modulatory signaling (TGF-β2), translational capacity (RRP15), secretory and trafficking processes (CNIH4), and cytoskeletal or transport organization (FEZ2). These loci represent priorities for fine-mapping and functional validation to further resolve the regulatory architecture underlying variation in myofiber hypertrophy in rainbow trout.

### Functional annotation of SNPs associated with myofiber CSA

A total of 117 SNPs were significantly associated with myofiber CSA across the genome. All SNPs were located on chromosome 2, indicating that most significant associations localized to a single genomic region consistent with a significant QTL (Table 2).

**Table 2.** Summary of SNPs significantly associated with myofiber cross-sectional area (CSA), including predicted functional impact categories and genomic distribution by variant type and region.

| Impact | Count | Percent |
| --- | --- | --- |
| Low | 6 | 1.46 |
| Modifier | 405 | 98.54 |

| Type | Count | Percent |
| --- | --- | --- |
| 3_prime_UTR_variant | 21 | 5.09 |
| 5_prime_UTR_variant | 4 | 0.97 |
| Downstream_gene_variant | 98 | 23.73 |
| Intergenic_region | 30 | 7.25 |
| Intron_variant | 221 | 53.51 |
| Splice_region_variant | 2 | 0.48 |
| Synonymous_variant | 4 | 0.97 |
| Upstream_gene_variant | 33 | 7.99 |

| Region | Count | Percent |
| --- | --- | --- |
| Downstream | 98 | 23.84 |
| Exon | 4 | 0.97 |
| Intergenic | 30 | 7.30 |
| Intron | 219 | 53.29 |
| Splice_Site_Region | 2 | 0.49 |
| Upstream | 33 | 8.03 |
| Utr_3_Prime | 21 | 5.11 |
| Utr_5_Prime | 4 | 0.97 |

Functional impact annotation showed that most SNP effects were classified as modifiers (405; 98.54%), with fewer low-impact (6; 1.46%) effects (Table 2). Consistent with the predominance of modifier effects, regional annotation indicated that most SNPs were located in noncoding regions, including introns (219; 53.285%), intergenic regions (30; 7.299%), and downstream regions (98; 23.844%), with additional variants in upstream regions (33; 8.029%). A smaller fraction of SNPs mapped to the 3′ UTR (21; 5.109%) and 5′ UTR (4; 0.973%), while relatively few were annotated to exons (4; 0.973%), or splice-site regions (2; 0.487%). Since SnpEff assigns non-mutually exclusive functional annotations, a single SNP may be classified into multiple categories simultaneously (e.g., both an intronic and upstream gene variant), resulting in annotation counts that exceed the total number of significant SNPs.

These patterns are consistent with the architecture commonly observed for complex traits, where trait-associated variants are enriched in noncoding regions and are more likely to act through gene regulation than by directly altering protein sequence. In particular, intronic, intergenic, and gene-flanking variants frequently overlap regulatory DNA and may influence promoter or enhancer activity, transcription factor occupancy, or chromatin state, thereby modulating expression of genes relevant to muscle growth and remodeling [36, 37]. The predominance of modifier-class annotations therefore supports a regulatory basis for myofiber CSA variation, while the presence of a concentrated signal on chromosome 2 indicates that one or more loci within the QTL interval contribute disproportionately to the phenotype.

Variant-type annotation supported the same pattern, with most variants classified as intron variants (221; 53.511%), intergenic region variants (30; 7.246%), and downstream gene variants (98; 23.729%), followed by upstream gene variants (33; 7.99%) and UTR-associated variants (3′ UTR variants: 21; 5.085% and 5′ UTR variants: 4; 0.969% (Table 2). Protein-altering annotations were uncommon, including synonymous variants (4; 0.969%), and splice-proximal variation was similarly limited (splice region variants: 2; 0.484%) (Table 2).

Overall, the functional annotation indicates that CSA-associated SNPs are dominated by noncoding, modifier-type variants, with most mapping to intronic, intergenic, and gene-flanking regions, and relatively few mapping to splice-site categories. This distribution, together with the strong clustering of associations on chromosome 2, is consistent with a significant QTL embedded within a broader background of regulatory variation, a pattern reported for many quantitative growth and musculoskeletal traits [36, 38].

### Functional annotation of SNPs associated with myofiber CSA at the *TGF-β2* locus

As shown in Table 3, significant SNPs associated with myofiber CSA at the *TGF-β2* locus are predominantly located within regions of regulatory function based on chromatin-state annotations, with the largest counts observed in enhancer-associated categories [18, 39]. Specifically, SNPs were enriched in weak enhancers (n = 6) and strong enhancers (n = 6), with additional variants in genic enhancers (n = 4) and a medium enhancer (n = 1), indicating that most associations at this locus map to regions linked to transcriptional regulation. Additional SNPs were identified in the 5′ UTR (n = 2), active transcription sites (n = 3), and transcription start flanking regions (n = 1).

**Table 3.**
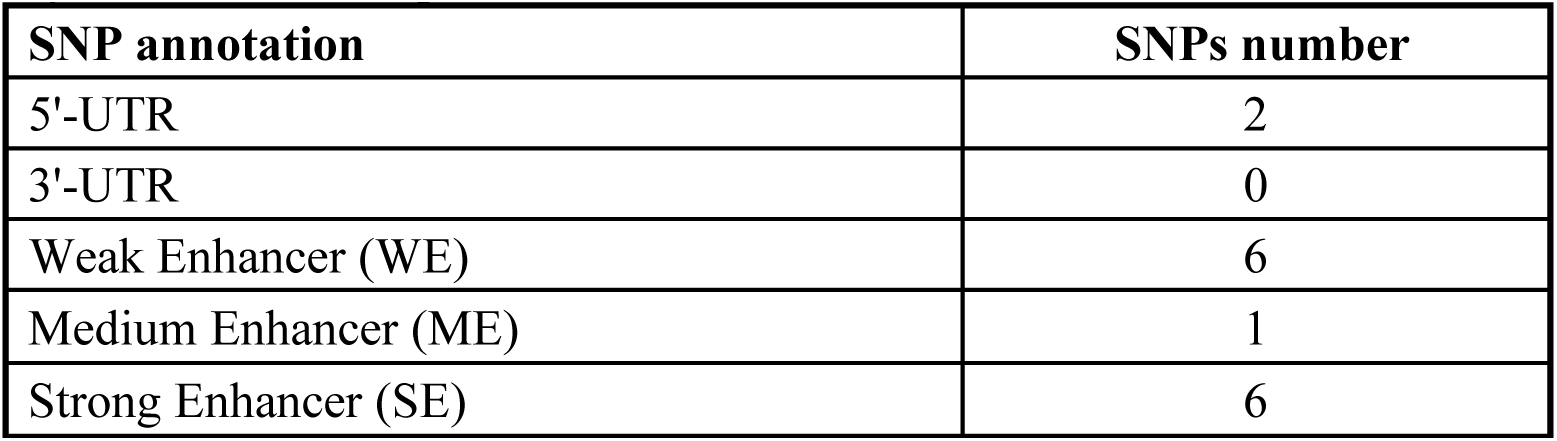
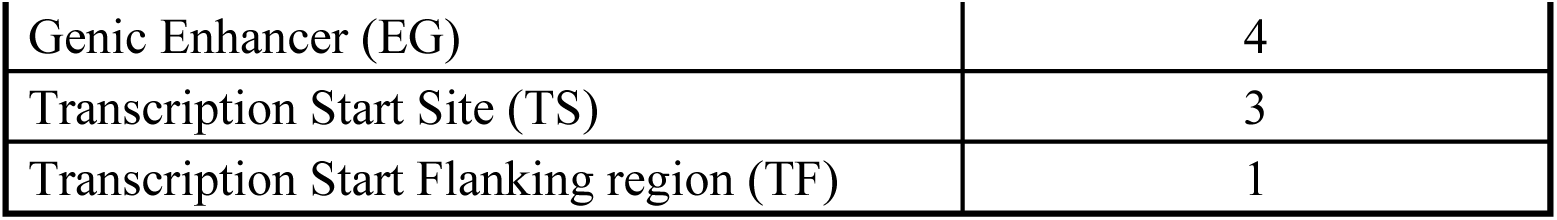
Distribution of significant SNPs across functional regions of the TGF-β2 gene, highlighting enrichment in regulatory annotations such as enhancer elements, untranslated regions, and transcription-associated sites.

| SNP annotation | SNPs number |
| --- | --- |
| 5'-UTR | 2 |
| 3'-UTR | 0 |
| Weak Enhancer (WE) | 6 |
| Medium Enhancer (ME) | 1 |
| Strong Enhancer (SE) | 6 |
| Genic Enhancer (EG) | 4 |
| Transcription Start Site (TS) | 3 |
| Transcription Start Flanking region (TF) | 1 |

Notably, consistent with IGV visualization, SNPs show overlapping localization with strong and genic enhancer elements as well as transcription start site–associated regions in white muscle, whereas this pattern is less consistent across other tissues. Although *TGF-β2* transcripts were not differentially expressed and SNPs were not concentrated within coding regions, variants at this locus colocalize with regulatory features including enhancer elements, transcription factor–associated regions, and untranslated regions. Together, these findings indicate that genetic variation at this locus is concentrated within regulatory architecture rather than protein-coding sequence.

The enrichment of associated variants in enhancer and UTR annotations is consistent with the established role of noncoding regulation in controlling gene expression and with evidence that trait-associated variants often localize to regulatory DNA rather than coding sequence [36–38].

*TGF-β2* is part of the TGF-β signaling network, which regulates cell proliferation, differentiation, and extracellular matrix remodeling, processes relevant to skeletal muscle growth and tissue architecture. Regulatory variation at *TGF-β2* therefore provides a plausible basis for altering signaling or expression programs that influence muscle fiber size, particularly when associated SNPs localize to enhancer and UTR annotations (Table 3), which are commonly implicated in transcriptional and post-transcriptional control [40].

Evidence from diverse vertebrate models supports a role for *TGF-β2* in muscle growth and remodeling. In neonatal rats, *TGF-β2* abundance in skeletal muscle has been reported to correlate positively with the fractional rate of muscle protein synthesis during early postnatal development, consistent with growth-stage–dependent involvement in protein accretion [41]. In fish, *TGF-β2* expression is dynamically regulated during muscle growth resumption and satellite cell differentiation, supporting its association with muscle growth and regenerative programs [31]. In chicken embryos, temporal expression patterns of *TGF-β2* transcripts during development have been linked to embryonic myogenesis, consistent with roles in early muscle formation [42]. In addition, TGF-β signaling is closely integrated with extracellular matrix regulation, and modulation of matrix-associated programs can influence cellular adhesion, migration, and differentiation—processes that shape muscle tissue remodeling and structure [43].

Together, the *TGF-β2* locus annotations (Table 3), combined with prior evidence for *TGF-β2* involvement in muscle growth-related processes across species, support a predominantly noncoding regulatory architecture for CSA-associated variation in this region [31].

### INDEL Genotype Variation Associated with Myofiber Cross-Sectional Area

To further refine genetic associations within the QTL region, we evaluated the relationship between indel variants and myofiber CSA. Association testing identified an indel (2_43212632_T_TGAGAGAGAGAGA) near the TGF-β2 locus that was significantly associated with the phenotype. The association between indel genotype and myofiber CSA was tested using an additive linear regression model that included genetic family as a covariate. A negative association was observed (INDEL β = −1,995, INDEL p = 2.67 × 10⁻⁴; model R² = 0.47, model p = 5.79 × 10⁻⁴), indicating that carriers of the alternative allele exhibited reduced myofiber CSA (Figure 5).

**Figure 4.**
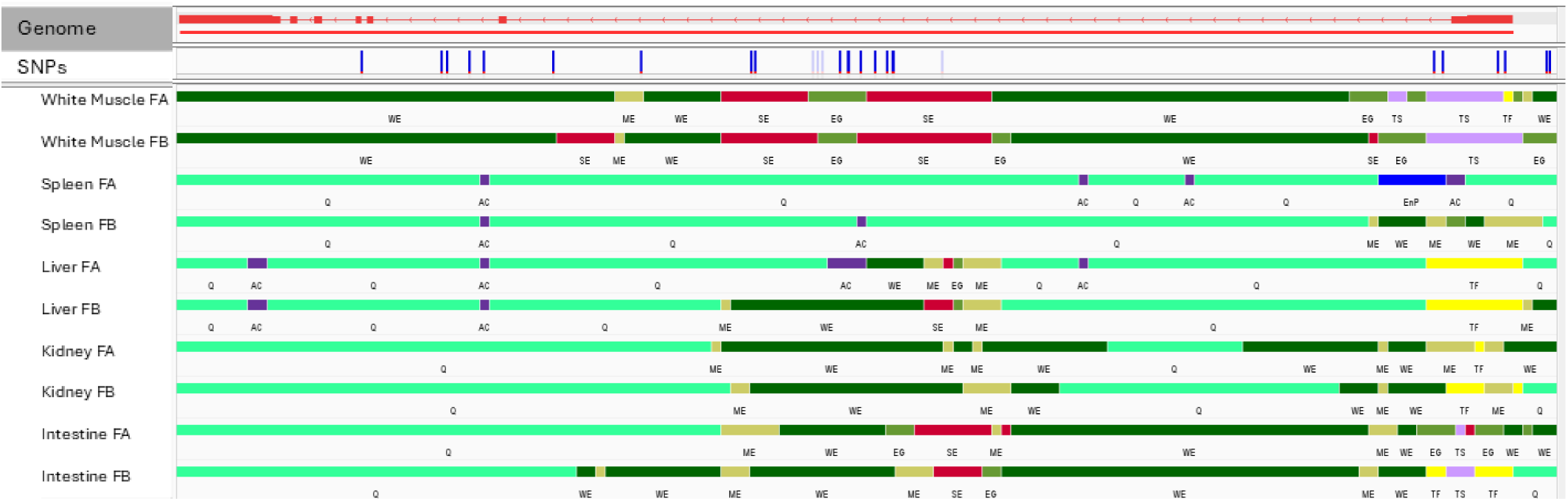
Integrated Genome Viewer (IGV) visualization of chromatin states at the TGF-β2 locus. SNPs are indicated by blue vertical lines across the gene region. A total of 21 SNPs were identified within the gene body, colocalizing with regulatory elements. Notably, in white muscle, SNPs consistently overlap strong enhancers (SE), genic enhancers (EG), weak enhancers (WE), and transcription start sites (TS), whereas this pattern is less consistent across other tissues. Chromatin state tracks are color-coded as shown.

**Figure 5.**
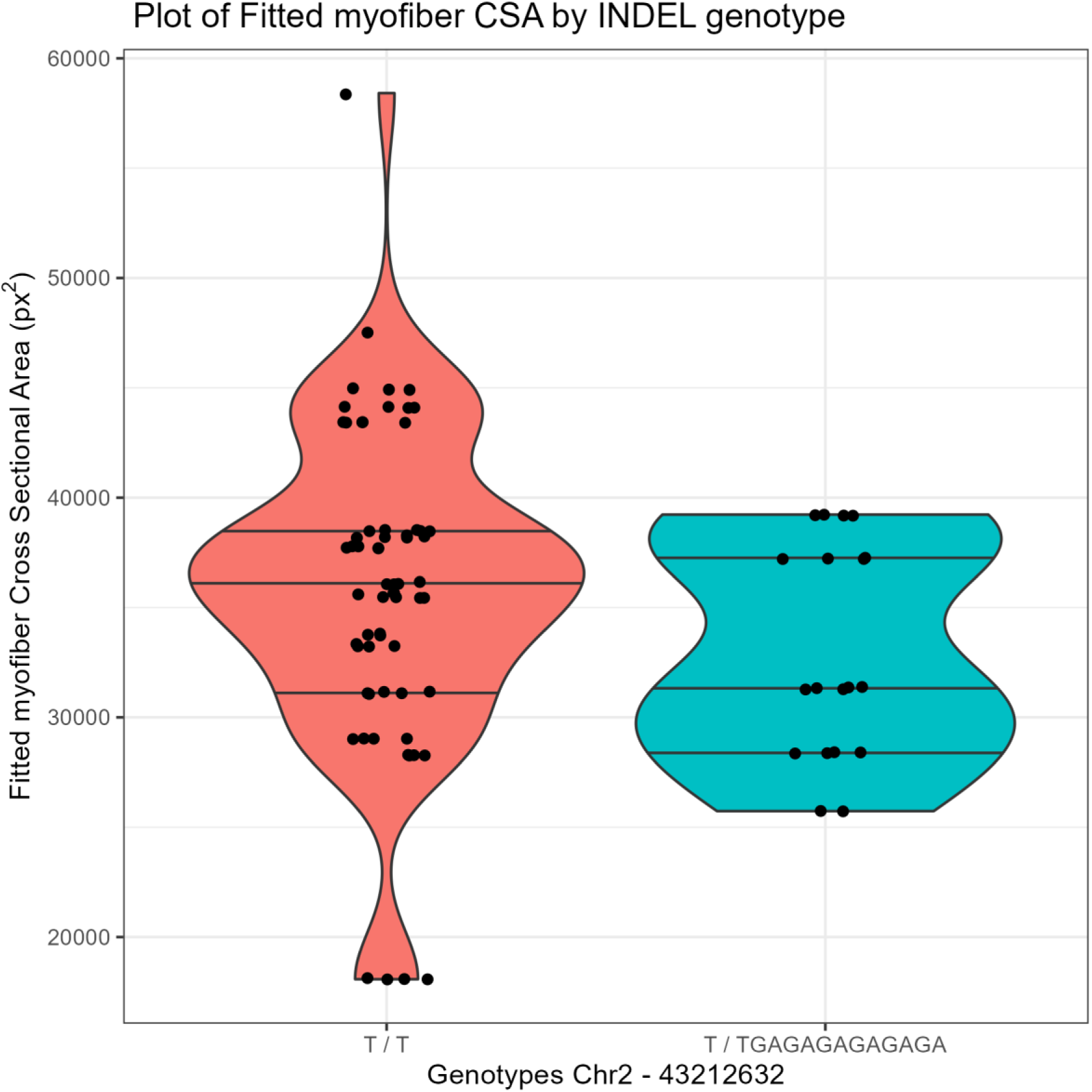
Distribution of regression-fitted myofiber cross-sectional area (CSA) across indel genotype classes at the TGF-β2 locus (2_43212632_T_TGAGAGAGAGAGA) for reference homozygous (T/T; n = 65) and heterozygous (T/ALT; n = 19) individuals. Horizontal lines represent the median and interquartile range. Carriers of the alternative allele exhibited significantly reduced myofiber CSA (β = −1,995; p = 2.67 × 10⁻⁴; model R² = 0.47).

Genotype distribution included 65 individuals with the reference genotype (T/T), 19 heterozygous individuals, and one homozygous alternative individual. Due to the low frequency of the homozygous alternative genotype, this individual was excluded from the analysis. Therefore, the reported association primarily reflects differences between reference homozygous and heterozygous individuals.

### Differential Gene Expression Associated with Muscle Phenotype Variation

To investigate molecular mechanisms underlying variation in muscle cellularity and related growth traits, transcriptome profiling was performed using individuals representing the highest and lowest quartiles of the myofiber CSA distribution. This design enabled contrasts between phenotypic extremes.

Differential expression analysis identified 73 genes with significant expression differences between the high and low myofiber CSA groups (FDR-adjusted P ≤ 0.05), spanning multiple biological pathways. Differentially expressed genes with roles relevant to muscle function are summarized in Table 4, and the complete list is provided in Supplementary File 2.

**Table 4.**
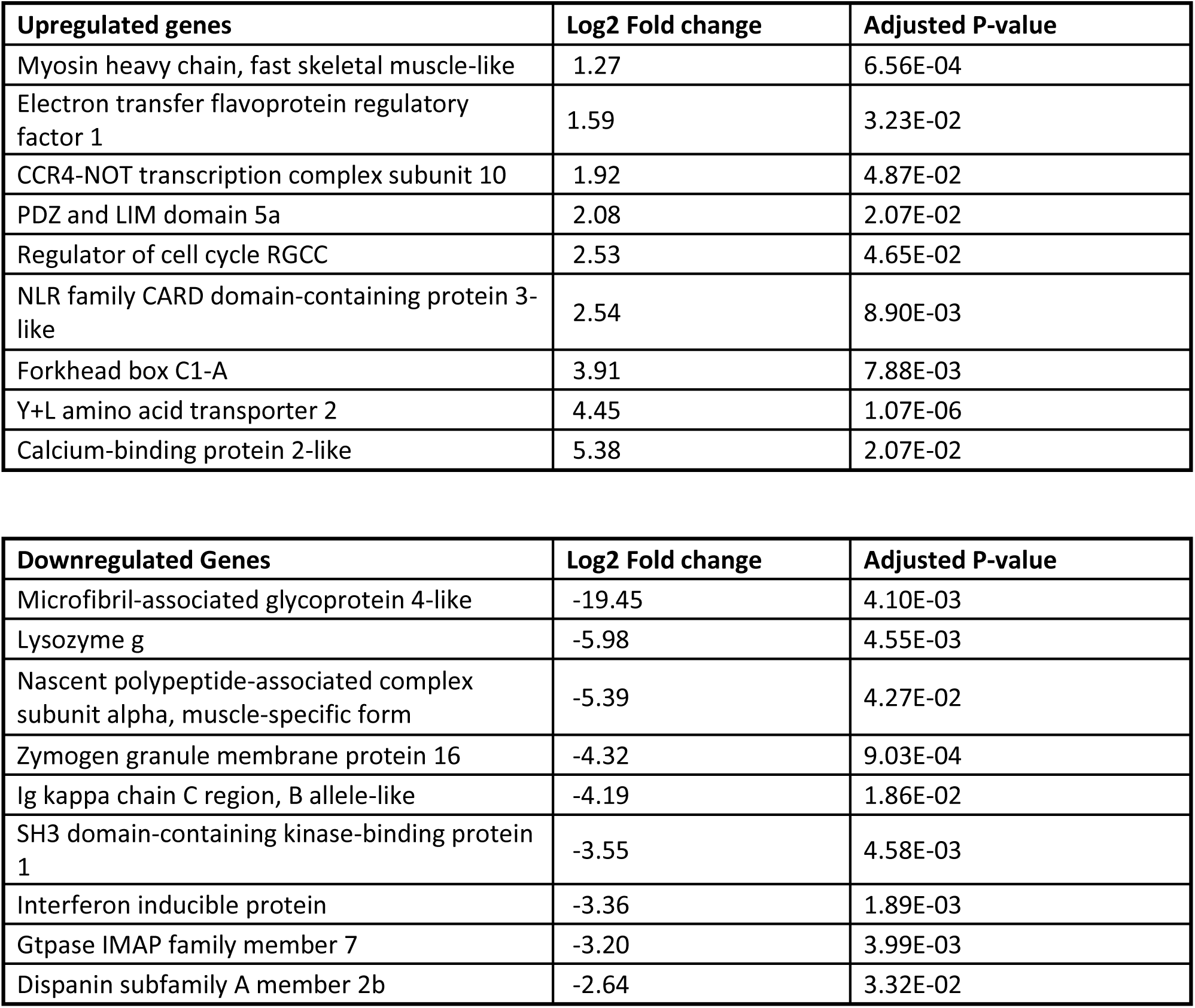
Selected differentially expressed genes between high and low myofiber CSA groups. Upregulated genes show higher expression in the high CSA group, and downregulated genes show lower expression in the high CSA group (FDR-adjusted P < 0.05).

| Upregulated genes | Log2 Fold change | Adjusted P-value |
| --- | --- | --- |
| Myosin heavy chain, fast skeletal muscle-like | 1.27 | 6.56E-04 |
| Electron transfer flavoprotein regulatory factor 1 | 1.59 | 3.23E-02 |
| CCR4-NOT transcription complex subunit 10 | 1.92 | 4.87E-02 |
| PDZ and LIM domain 5a | 2.08 | 2.07E-02 |
| Regulator of cell cycle RGCC | 2.53 | 4.65E-02 |
| NLR family CARD domain-containing protein 3-like | 2.54 | 8.90E-03 |
| Forkhead box C1-A | 3.91 | 7.88E-03 |
| Y+L amino acid transporter 2 | 4.45 | 1.07E-06 |
| Calcium-binding protein 2-like | 5.38 | 2.07E-02 |

| Downregulated Genes | Log2 Fold change | Adjusted P-value |
| --- | --- | --- |
| Microfibril-associated glycoprotein 4-like | -19.45 | 4.10E-03 |
| Lysozyme g | -5.98 | 4.55E-03 |
| Nascent polypeptide-associated complex subunit alpha, muscle-specific form | -5.39 | 4.27E-02 |
| Zymogen granule membrane protein 16 | -4.32 | 9.03E-04 |
| Ig kappa chain C region, B allele-like | -4.19 | 1.86E-02 |
| SH3 domain-containing kinase-binding protein 1 | -3.55 | 4.58E-03 |
| Interferon inducible protein | -3.36 | 1.89E-03 |
| Gtpase IMAP family member 7 | -3.20 | 3.99E-03 |
| Dispanin subfamily A member 2b | -2.64 | 3.32E-02 |

### High Myofiber CSA Muscle Exhibits a Transcriptional Program Optimized for Hypertrophic Growth

Skeletal muscle constitutes the major proportion of body mass in rainbow trout, and post-embryonic growth reflects coordinated regulation of muscle cellularity and accumulation of myofibrillar proteins [44]. In this study, genes annotated as myosin heavy chain, fast skeletal muscle-like *(MyHC)* were upregulated in the high myofiber CSA group. This finding is consistent with increased expression of a core sarcomeric component in individuals exhibiting greater muscle myofiber CSA.

MyHC is the principal motor protein of the thick filament and is central to cross-bridge cycling and force generation during contraction [45–47]. In fish, *MyHC* transcript abundance has been shown to track muscle protein accretion, supporting its use as a marker of myofibrillar growth state [46]. Taken together, the upregulation of fast skeletal muscle-like *MyHC* in the high myofiber CSA group indicates that differences in muscle phenotype are accompanied by differential expression of contractile machinery, consistent with a greater investment in myofibrillar components in fish with larger myofiber CSA. The results support a conservative conclusion that variation in myofiber CSA is associated with altered expression of genes directly involved in sarcomere structure and function, linking histological differences in muscle phenotype to transcriptional differences in the contractile apparatus.

We report PDZ and LIM domain 5 (*PDLIM5*) as upregulated in the high myofiber CSA group. PDLIM5 is a PDZ-LIM family protein that functions as an actin-associated scaffold, positioning binding partners at cytoskeletal sites and linking cytoskeletal organization with regulatory protein interactions [48, 49]. PDZ-LIM proteins have also been reported to participate in nucleocytoplasmic interactions, including binding partners with nuclear roles and anchoring them at cytoplasmic sites [49]. In addition, experimental work in chicken skeletal muscle satellite cells has shown that PDLIM5 can influence proliferation and differentiation through p38-MAPK–associated signaling [50]. While these cellular processes were not directly evaluated here, these studies provide context for the association between *PDLIM5* upregulation and variation in muscle cellularity in rainbow trout.

Calcium-binding protein 2-like (*CABP2-like*) was upregulated in the high myofiber CSA group, indicating differential expression of a Ca^2+^-binding regulator in individuals with larger muscle cellular area. *CABP2* belongs to the calcium-binding protein (*CaBP)* family, a set of calmodulin-like Ca^2+^ sensors that share structural similarity with calmodulin and participate in Ca^2+^-dependent signal modulation [51]. Gene function summaries for *CABP2* further describe this family as capable of interacting with calmodulin-linked effector pathways (e.g., *CaMKII* and calcineurin), although these interactions have not been specifically established in rainbow trout skeletal muscle [52].

Ca^2*+*^ handling is central to excitation–contraction coupling, where depolarization triggers Ca^2*+*^ release from the sarcoplasmic reticulum and Ca^2*+*^ binding to troponin activates actin–myosin interaction and force generation [53]. Because Ca^2*+*^ cycling governs contraction–relaxation kinetics and supports muscle plasticity, differential expression of Ca^2*+*^ regulatory genes can accompany differences in muscle functional state and remodeling. [54]. Consistent with this context, studies in other vertebrates emphasize that maintaining intracellular Ca^2*+*^ homeostasis is closely tied to preservation of skeletal muscle integrity under challenging physiological conditions [55].

Taken together, the upregulation of *CABP2-like* in the high myofiber CSA group suggests that variation in muscle cellularity is accompanied by transcriptional differences in Ca^2*+*^-associated regulation and excitation–contraction biology, without implying a specific causal mechanism or pathway activation in the absence of functional measurements.

Forkhead box (FOX) transcription factors are pleiotropic regulators of gene expression with established roles in cell-cycle control, proliferation, differentiation, and developmental patterning [56]. In teleosts, *FOXc1a* has been implicated in early somitogenesis: loss of *FOXc1a* function in zebrafish disrupts somite formation and is associated with altered Notch and FGF signaling during somite differentiation [57]. Consistent with a conserved role for *FOXc* factors in paraxial mesoderm development, mouse studies show that combined *FOXc1/FOXc2* dosage influences somite maturation and axial skeleton development [58]. In addition to embryonic patterning, *FOXc1* has been linked to myogenic regulation: a circCPE/miR-138/FOXC1 regulatory network was reported to modulate bovine myoblast development, supporting a role for *FOXc1* in muscle cell proliferation and differentiation programs [59]. Given that *FOXc1a* is upregulated in our high muscle yield and high cell-size line, these prior findings support the interpretation that variation in muscle phenotypes may be accompanied by transcriptional differences in developmental and myogenic regulatory pathways.

Electron transfer flavoprotein regulatory factor 1 (*ETFR1*; also annotated as *ETFRF1/LYRM5* in vertebrates) was upregulated in the high muscle yield genetic line and the high cell-size phenotype. ETFRF1 is a mitochondrial LYR-motif protein implicated in the respiratory electron transport chain through regulation of electron transfer flavoprotein (ETF), a central electron shuttle that accepts electrons from multiple mitochondrial dehydrogenases and transfers them to the main respiratory chain via ETF:ubiquinone oxidoreductase (ETF-QO/ETFDH) [60]. In this context, elevated *ETFR1/ETFRF1* expression is consistent with transcriptional differences in mitochondrial bioenergetic regulation, which is relevant because ATP supply supports energy-demanding processes associated with muscle growth, including protein synthesis and contractile activity.

CCR4-NOT transcription complex subunit 10 (*CNOT10)* was upregulated in the high cell-size phenotype._The *CCR4-NOT* complex is a multisubunit regulator present in all eukaryotes that coordinates gene expression across multiple stages, from nuclear transcription-associated processes to cytoplasmic translation and mRNA decay [61]. In the cytoplasm, it influences mRNA translatability, contributes to co-translational quality control, and can promote translational repression as well as decapping and deadenylation-driven mRNA turnover [61]. In metazoans, *CNOT10* is a conserved subunit that, together with *CNOT1*1, binds the N-terminal region of CNOT1 to form a stable NOT10:NOT11 module that contributes to CCR4-NOT architecture and function [62, 63]. In our dataset, *CNOT10* was upregulated in the high muscle yield and high cell-size line, consistent with transcriptional differences in regulatory programs that shape translational output and mRNA turnover.

Y+L amino acid transporter 2 was upregulated in fish with high myofiber CSA. The Y+L amino acid transporter 2 (*SLC7A6*) is an obligatory exchanger that mediates the uptake of cationic amino acids, such as L-arginine, in exchange for the efflux of neutral amino acids like L-glutamine, with transport supported by Na+ gradients [64, 65]. In this study, the upregulation of *SLC7A6* in fish with high myofiber CSA is consistent with the established role of solute carrier transporters in regulating cellular amino-acid flux and proteostasis [66, 67]. Increased *SLC7A6* activity would be expected to enhance the availability of L-arginine, the precursor for nitric oxide (NO), and thereby support NO-dependent regulation of skeletal muscle blood flow and activation of myogenic satellite cells that contribute to fiber hypertrophy [68, 69]. Furthermore, by modulating intracellular pools of essential amino acids, including L-leucine, *SLC7A6* could contribute to activation of the mTORC1 pathway, a central regulator of muscle protein synthesis [67, 70]. Consequently, elevated *SLC7A6* expression is compatible with a transcriptional signature of enhanced nutrient sensing and anabolic signaling that underpins muscle enlargement.

### Low myofiber CSA muscle shows remodeling and ECM-immune activity

Nascent polypeptide-associated complex subunit alpha, muscle-specific form was upregulated in the low myofiber CSA. The skeletal and cardiac muscle-specific isoform of *NACA* (*skNAC)* is generated by differential inclusion of a large proline-rich exon, converting the ubiquitous αNAC into a muscle-restricted transcription factor [71]. *skNAC* expression is induced during myogenic differentiation and has been linked to muscle repair and regeneration programs in vivo [72]. Consistent with this role, targeted disruption of skNAC in mice results in reduced postnatal skeletal muscle growth and impaired regenerative capacity following injury [73]. In zebrafish, *skNAC* is required for normal myofibril organization during embryonic muscle development [74].

In our dataset, *skNAC* was upregulated in the low myofiber CSA (low muscle weight) group. One interpretation is that low myofiber CSA muscle may reflect relatively greater remodeling or regeneration-associated activity, with higher *skNAC* expression marking ongoing differentiation-related transcriptional programs.

The low myofiber CSA phenotype showed coordinated upregulation of extracellular matrix (ECM) and immune-related genes, consistent with a muscle environment that is more remodeling-and inflammation-associated than hypertrophy-associated. *MFAP2 (MAGP-1)* and *MFAP4*-like encode microfibril-associated proteins linked to fibrillin-rich microfibrils, and MAGP proteins can bind active TGF-β/BMP ligands and modulate their signaling context [75]. In teleosts, *MFAP4* is also strongly associated with the macrophage lineage and has been implicated in host defense, providing a plausible molecular link between ECM remodeling and innate immune activity [76]. Consistent with this, low myofiber CSA tissue also showed higher expression of immune markers (e.g., lysozyme and interferon-inducible genes), suggesting elevated immune activation. Prolonged or excessive inflammatory signaling is well documented to impair muscle recovery and promote wasting in vertebrates [77]. In rainbow trout, stress physiology can suppress growth through endocrine and feeding-related mechanisms, including reduced circulating growth hormone under acute stress and inhibition of appetite and nutrient-sensing pathways under stress [78]. Collectively, these patterns support an association between increased ECM/immune activity and reduced muscle accretion in the low myofiber CSA group, while remaining correlative.

Together, these results support a model of two distinct muscle states associated with myofiber CSA: (1) a hypertrophy-associated, anabolic state characterized by increased expression of genes involved in contractile function, cytoskeletal organization, and translational capacity, and (2) a remodeling-associated state marked by elevated extracellular matrix and immune-related gene expression.

## CONCLUSION

This study establishes myofiber CSA as a key histological bridge between genetic variation and whole-animal growth traits in rainbow trout, with larger myofiber CSA associated with higher body weight, muscle weight, visceral weight, and body length. By integrating histology, GWAS, and transcriptomics, a QTL was resolved on chromosome 2 (Omy2), where 117 significant SNPs cluster within a 103.8 Mb window overlapping regulatory regions near *TGF-β2*, and additional growth-related genes.

Although *TGF-β* ligand genes were not differentially expressed, the enrichment of variants in enhancers and untranslated regions at this locus suggests that muscle growth potential may be tuned through changes in signaling sensitivity or ligand bioavailability rather than through altered transcript abundance. Transcriptomic profiling further supports a model of contrasting muscle states: high myofiber CSA fish display an anabolic program with upregulation of contractile machinery *(MyHC)*, cytoskeletal scaffolds *(PDLIM5)*, and signaling regulators *(CaBP2*), whereas low myofiber CSA fish upregulate extracellular matrix and immune genes (e.g., *MAGP-1*–associated pathways), indicative of a more dynamic remodeling- and defense-oriented muscle state. Together, these results extend phenotypic associations by providing a molecular framework linking a significant QTL enriched for regulatory variation to distinct transcriptional states associated with muscle fiber hypertrophy. This study highlights the role of noncoding regulatory mechanisms in shaping muscle cellularity and growth-related traits in rainbow trout.

## DECLARATIONS

### Ethics approval and consent to participate

All husbandry and experimental procedures were approved by the Institutional Animal Care and Use Committee (IACUC) at the University of Maryland, College Park (protocol #1593175-6).

### Consent for publication

Not applicable

### Availability of data and materials

Transcriptomic data is available in BioProject accession number: PRJNA974903 https://www.ncbi.nlm.nih.gov/bioproject/PRJNA974903

All pertinent scripts are available on GitHub: https://github.com/akashr-umd/Salem-Lab--Muscle-Histology-Project

### Competing interests

The authors declare that they have no competing interests.

### Funding

This study was supported by competitive grants No. 2023-67015-39742 from the United States Department of Agriculture, National Institute of Food and Agriculture.

### Authors’ contributions

AR performed histology image collection, data collection, and drafted the manuscript. GR, RA, and AA conducted data analysis, contributed to figure production, and edited the manuscript. TL and MS performed final edits to the manuscript. MS conceived and designed the experiments. All authors read and approved the final manuscript.

## Acknowledgements

Not applicable

## Supplementary Files

Supplementary File 1. contains phenotypic data of individual fish CSA, body weight, muscle weight, visceral weight, body length, and genetic lines.

Supplementary File 2. contains differentially expressed genes and fold changes for RNA sequencing data.

Supplementary Figure 1. contains QQ plot showing genomic inflation (λGC) calculated to be 1.02, suggesting there is no bias or inflation in our analysis.

## Notes

https://www.ncbi.nlm.nih.gov/bioproject/PRJNA974903

https://github.com/akashr-umd/Salem-Lab--Muscle-Histology-Project

### Competing Interest Statement

The authors have declared no competing interest.

