## Supplementary figures and images for "QTL spanning the TGF-β2 locus is associated with muscle fiber hypertrophy in rainbow trout"

### Supplementary Figure 1

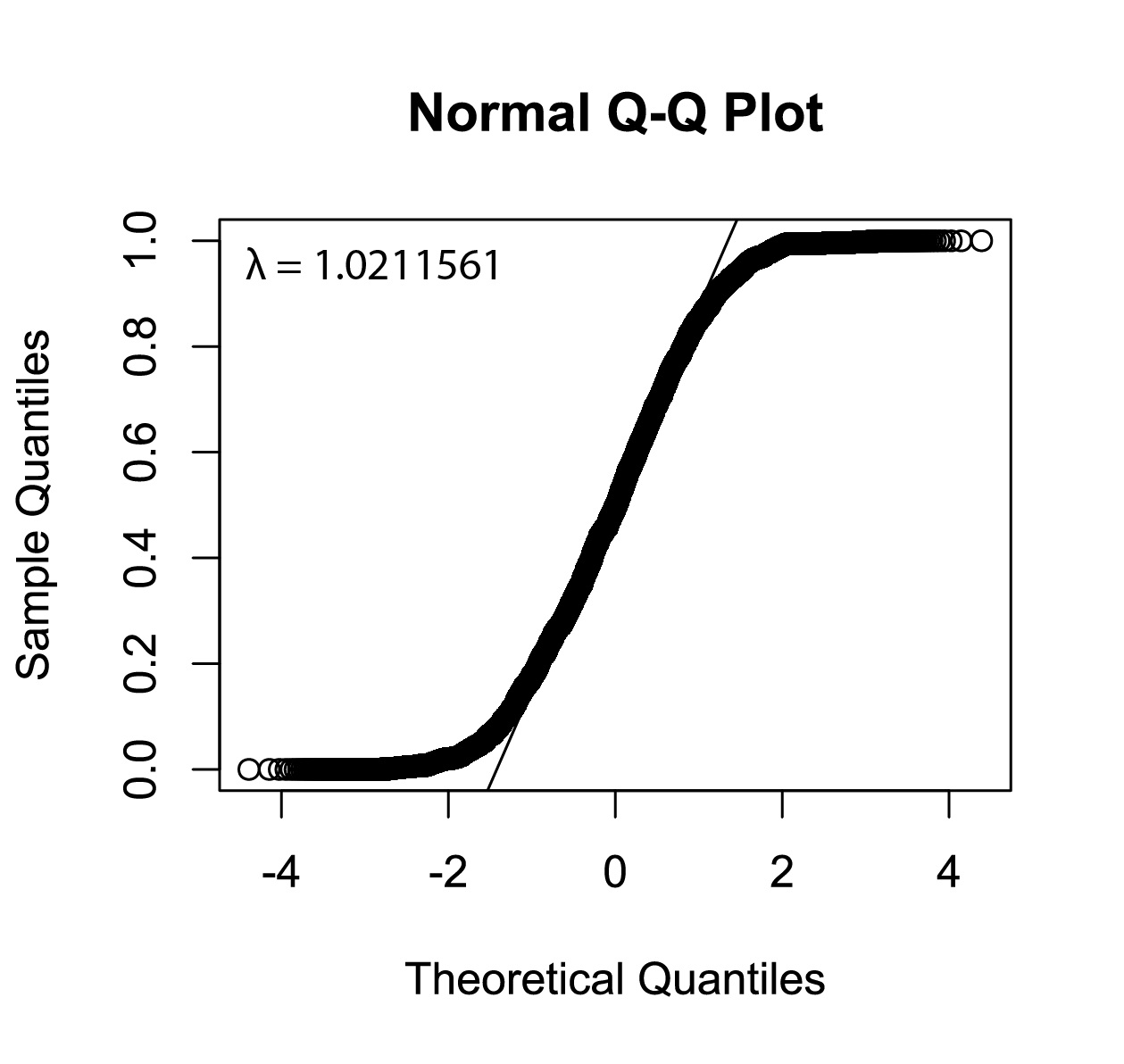
